# Pharmacological inhibition of Fms-like kinase 3 (FLT3) promotes neuronal maturation and suppresses seizure in a mouse model

**DOI:** 10.1101/2024.10.02.616380

**Authors:** Kate Cruite, Madeline Moore, Michael D. Gallagher, Emanuel M. Coleman, Yuqin Yin, Shuqi Lin, Ellie Shahbo, Tenzin Lungjangwa, Volker Hovestadt, Rudolf Jaenisch, Jamie Maguire, Xin Tang

**Affiliations:** Department of Neurosurgery, Boston Children’s Hospital, Boston, MA, USA; Whitehead Institute for Biomedical Research, Cambridge, MA USA; Tufts University School of Medicine, Boston, MA, USA; Dana Farber Cancer Institute, Boston, MA, USA; Harvard Medical School, Boston, MA, US

## Abstract

Fms-like tyrosine kinase 3 (FLT3) is a receptor tyrosine kinase predominantly expressed in blood and brain cells. While the FLT3 signaling pathway has been extensively studied in blood cell development and leukemia, its role in the brain remains largely unexplored. Through our group’s previous high-throughput drug screening work unexpectedly found that several small molecule FLT3 inhibitors (FLT3i), including KW-2449 and Sunitinib, enhance expression of the gene encoding chloride transporter KCC2 in neurons. KCC2 is crucial for brain development and function, and its dysregulation is linked to many brain diseases. These findings suggest previously unrecognized roles of FLT3 signaling in brain health and disease that have yet to be systematically studied. In this study, we utilized a functional genomics approach to investigate the transcriptomic changes induced by pharmacological inhibition of the FLT3 pathway in brain cells, including cultured primary mouse neurons, a human stem cell-derived neuronal model of Rett syndrome (RTT), and human stem cell-derived microglia cultures. Our results show that treating human or mouse neurons with FLT3i drugs significantly upregulates genes crucial for brain development while downregulating genes linked to neuroinflammation. In contrast, FLT3i treatment of human microglia, which do not express FLT3, has no effect on their gene expression, highlighting the cell type-specific roles of FLT3 signaling in the brain. To further understand how FLT3 signaling regulates the expression of neuronal maturation genes such as KCC2, we conducted a curated CRISPR screen that identified a number of transcription factors involved in FLT3i-mediated KCC2 activation in neurons. The mRNA and protein levels of several neurodevelopmental disorder (NDD) risk genes are significantly upregulated in FLT3i-treated neurons, indicating potential therapeutic applications of FLT3i in rescuing underexpression and/or haploinsufficiency of disease-associated genes. In our *in vivo* studies, we evaluated the efficacy of the FLT3i drug KW-2449 in mice, demonstrating that it can effectively cross the blood-brain barrier, induce KCC2 protein expression for up to 24 hours after a single injection, and reduces seizure activity in a chemoconvulsant-induced mouse model of temporal lobe epilepsy. Collectively, our findings uncover previously unrecognized roles of neuron-specific FLT3 signaling in promoting neuronal maturation and reducing neuroinflammation. These results suggest that FLT3 kinase signaling regulates a transcriptional program vital for brain development and function, position it as a promising therapeutic target for NDD treatment.

## Introduction

Kinase signaling plays major roles in mediating signal transduction cascades in various cell types^*1*^. Drugs that target kinase signaling pathways and have been developed to combat diseases such as cancer and brain disorders^*2*^. The receptor tyrosine kinase FLT3 is activated by the FLT3 ligand, a pleiotropic cytokine abundant in blood and cerebrospinal fluid^*3, 4*^. Investigations of FLT3 kinase signaling have historically been focused on understanding its role in the proliferation and differentiation of blood cells, where somatic tandem duplication of the FLT3 juxtamembrane domain leads to overactivation of the FLT3 signaling pathway, causing leukemia^*5*^. In contrast, very little is known about the role of the FLT3 pathway in brain health and disease^*6*^. Global knockout of the *Flt3* gene in mice leads to reduced production of primitive hematopoietic progenitors in mice older than six months^*7*^, but does not cause any behavior or neurological deficits (International Mouse Phenotyping Consortium, MGI 95559). Missense and loss-of-function variants of the *FLT3* gene are widely present in the human population and are not associated with any known developmental defect or genetic disease (Genome Aggregation Database gnomAD constraint metrics score = 0.98 and 0.61, respectively). These results suggest that FLT3 may be a safe molecular target to modulate for therapeutic benefit.

Although the presence of FLT3 ligand (FL) and the FLT3 receptor tyrosine kinase proteins in the brain has been known for decades^*8, 9*^, few studies have directly investigate FLT3’s roles in brain development and functioning^*6*^. A recent transcriptome-wide association study reveals that the pathogenesis of Tourette syndrome, a neurodevelopmental and epileptic disorder, is associated with an upregulation of neuronal *FLT3* mRNA expression in the prefrontal cortex^*10*^. An increased level of FLT3 ligand in cerebrospinal fluid (CSF) has been identified as a biomarker associated with neurodegenerative disorders including Sjögren’s syndrome^*11*^, amyotrophic lateral sclerosis^*9*^, and Parkinson’s disease^*12*^. Excessive FLT3 signaling has also been revealed as a key regulator of neuropathic pain, a process strongly associated with dysregulation of neuronal chloride transporter KCC2 that results in hyperexcitability of sensory circuits^*13*^. Injection of the FLT3 ligand into the spinal cord causes sensory neuron overexcitation and chronic pain^*14*^, while inhibition of FLT3 signaling alleviates peripheral neuropathic pain in mice^*15*^. Taken together, these findings indicate that Flt3 signaling may play major roles in brain health and disease, although the mechanism remains elusive.

Small molecule drugs that inhibit FLT3 kinase pathway signaling (FLT3i) have been developed to treat leukemia and solid tumors. Among these are KW-2449, which was tested in acute myeloid leukemia (AML) clinical trial to treat FLT3 ITD leukemia; and Sunitinib which is in clinical use for the management of AML^*16*^. Our previous work aiming to screen for small molecule compounds that activate the expression of the *SLC12A5* gene, which encodes the neuron-specific potassium-chloride co-transporter KCC2, led to the unexpected discovery that a number of FLT3i drugs, including KW-2449 and Sunitinib, stimulate the expression of KCC2 mRNA and protein in neurons^*17*^. KCC2 is a Cl^-^ transporter required for GABAergic inhibition in the brain.

Impairment in GABAergic inhibition is a fundamental mechanism underlying the hyperexcitability of neural circuits in epilepsy. Dysregulation of KCC2 due to genetic mutation, epigenetic expression silencing, and excessive protein degradation plays a pivotal role in regulating the initiation and recurrence of epilepsy^*18*^. Epilepsy is a serious condition associated with substantial individual and societal cost, affecting three million adults and 470,000 children in the US, with 1.2% of the total US population having had active epilepsy during their lifetime^*19*^. The involvement of KCC2 in epilepsy makes it a promising –yet underexplored-drug target for anti-seizure medications. More than one third of all epilepsy patients do not respond to any available anti-seizure medications^*20*^, indicating a critical unmet need for the development of novel therapeutic approaches. Since FLT3i drug stimulates KCC2 mRNA and protein production, it may represent a novel therapeutic avenue for anti-seizure medication development.

In our current study, we provide the first systematic characterization of the cellular specificity and gene expression changes induced by FLT3i treatment of brain cells. We find that FLT3i treatment stimulates a transcription program associated with the functional maturation of neurons, including expression of a number of NDD risk genes, while suppressing the expression of multiple neuroinflammation genes, in cultured mouse neurons and in human stem cell-derived neurons but not in microglia that do not express FLT3. Further, we have identified several novel FLT3i-responsive transcription factors that regulate KCC2 expression in neurons, and demonstrated the capability of the FLT3i drug KW-2449 to enter the brain, induce KCC2 protein expression *in vivo*, and exhibit anti-seizure efficacy in a mouse model of pharmacoresistant epilepsy. Through elucidating previously unknown mechanisms of FLT3 signaling in brain development and diseases, our results provide a foundation for further development of a novel class of FLT3i gene modulation drugs that can be tailored/matched to disease-relevant molecular signatures caused by NDD risk gene deficiencies to offer therapeutic benefit.

## Results

### FLT3i treatment stimulates the expression of neuronal maturation genes while suppressing neuroinflammation genes in cultured mouse neurons

Kinases transmit signals primarily through modulating the phosphorylation status of their substrates, which in turn can alter the expression level of target genes in cells^*21*^. To systematically unravel the genes regulated by FLT3 kinase signaling in the brain, we treated cultured primary mouse cortical neurons with KW-2449 or Sunitinib, two structurally distinct small molecule FLT3i compounds identified and validated from our previous drug screening work^*17*^, then performed mRNA sequencing (RNA-seq) to provide the first unbiased characterization of FLT3i-induced gene expression changes in neurons (**Fig. 1A**). Our results show that FLT3i drug treatment of cultured primary mouse neurons induces robust and reproducible gene expression changes (**Fig. 1B**). Compared to DMSO-treated control neurons, both KW-2449- and Sunitinib-treated mouse neurons exhibit a large number of differentially expressed genes (**Fig. 1C-D**). Interestingly, a strong positive correlation (R = 0.72) can be detected between the DMSO control-normalized expression level changes in individual genes responding to either KW-2449 or Sunitinib treatment (**Fig. 1E**), indicating the engagement of a common set of transcriptional programs in response to these two agents. We then performed differential gene expression analysis using the DESeq2 bioinformatics package, defining differentially expressed genes (DEGs) as having an adjusted p-value smaller than 0.05 and absolute value of the log2 fold-change greater than 1. Compared to DMSO-treated control neurons, KW-2449 treatment leads to ∼ 800 upregulated and ∼2000 downregulate DEGs, whereas Sunitinib induces ∼200 upregulated and ∼800 downregulated DEGs. Importantly, the KW-2449-induced and Sunitinib-induced DEG gene lists show substantial overlap (**Fig. 1E** insert). Such strong correlation in gene expression pattern induced by chemical compounds that are structurally distinct, and therefore have different off-target bindings, strongly suggest a common on-target mechanism of inhibiting FLT3 signaling.

**Figure 1:**
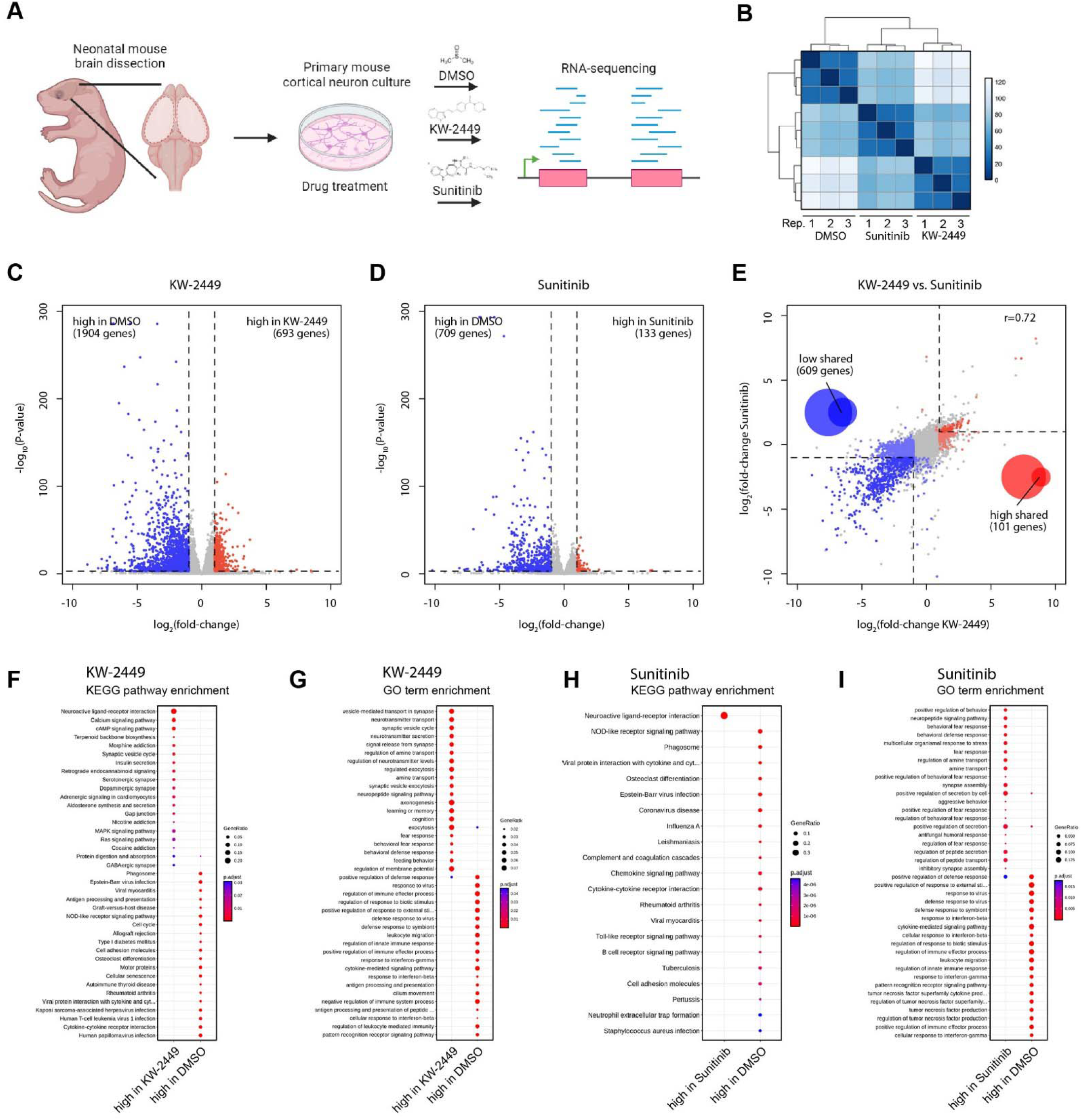
Treatment of cultured mouse neurons with FLT3 kinase inhibitors induce robust and reproducible gene expression changes. (**A**) Diagram showing the experimental workflow: (**B**) Correlational matrix dendrogram representation of RNA-seq results from different treatment conditions and repeats. (**C-D**) Volcano plots showing the differentially-expressed genes (DEGs) in KW-2449 or Sunitinib-treated mouse neurons. (**E**) Scatter plot illustrating the normalized Log_2_ fold change values of individual genes in response to KW-2449 or Sunitinib treatment. The Venn diagram insets display the significant overlap between DEGs under the two drug treatment conditions, highlighting shared and unique gene expression changes. (**F-I**) Gene Ontology (GO) and KEGG pathway analyses of DEGs induced by KW-2449 and Sunitinib reveal upregulation of synaptic genes and downregulation of genes associated with neuroinflammation.

We further analyzed the RNA-seq results through using Kyoto Encyclopedia of Genes and Genomes (KEGG) pathway analysis and Gene Ontology (GO) analysis of DEGs to identify molecular pathways and processes associated with FLT3i treatment. Our analysis revealed that both KW-2449 and Sunitinib treatment results in robust upregulation of gene sets associated with neuronal maturation, such as neuroactive ligand-receptor interaction, synaptic vesicle cycle, and inhibitory synapse assembly. At the same time, gene sets associated with inflammatory processes, such as cytokine-cytokine receptor interactions, regulation of immune effector processes, and response to interferon-gamma, are enriched in DEGs downregulated by FLT3i treatment (**Fig. 1F-I**). Such coordinated patterns of gene regulation indicate that FLT3 signaling modulates gene co-regulatory networks that play important roles in brain health regarding both neuronal functional maturation and neuroinflammation.

### FLT3i treatment elicits transcriptomic alterations in human neurons that are similar to those observed in mouse neurons

We next assessed the extent to which the transcriptomic changes induced by FLT3i treatment in brain cells is conserved across species. Our previous work demonstrated that treatment with FLT3i drug KW-2449 stimulates KCC2 expression and corrects electrophysiological abnormalities in an *MECP2* knockout human stem cell-derived neuronal model of Rett syndrome^*17*^. We conducted drug treatment followed by RNA-seq experiments showing that KW-2449 treatment of human RTT neurons induces reproducible gene expression changes in human neurons across three biological repeats with hundreds of DEGs (**Fig. 2A-C**). Interestingly, KEGG and GO analysis of the FLT3i-induced DEGs in mouse and human neurons reveal a substantial overlap in functional categories: the upregulated gene categories include neuroactive ligand-receptor interaction, calcium signaling, and synaptic vesicle cycling; while the downregulated gene categories include cytokine-cytokine receptor interaction (**Fig. 2D-E**). We further performed gene set enrichment analysis (GSEA) to systematically profile the FLT3i-induced changes in gene expression pattern. We detected enrichment in gene sets associated with neuronal maturation including neurotransmitter release, neuroactive ligand-receptor interaction, neurotransmitter receptors, synaptic vesicle cycle, and ion channels (**Fig. 3A**). On the other hand, FLT3i treatment of neurons deplete gene sets associated with neuroinflammation, including genes involved in the chemokines, interferon, TNF, and JAK-STAT signaling pathways (**Fig. 3B**). At the individual gene level, the FLT3i-induced gene expression changes show remarkable consistency across different drug treatment conditions (KW-2449 and Sunitinib) and in both human and mouse neurons, in both the up- and down-regulated gene sets (**Fig. 3B, D**). Our results indicate that the conserved FLT3i-responsive genes are functionally important and drive similar biological processes in both mouse and human neurons.

**Figure 2:**
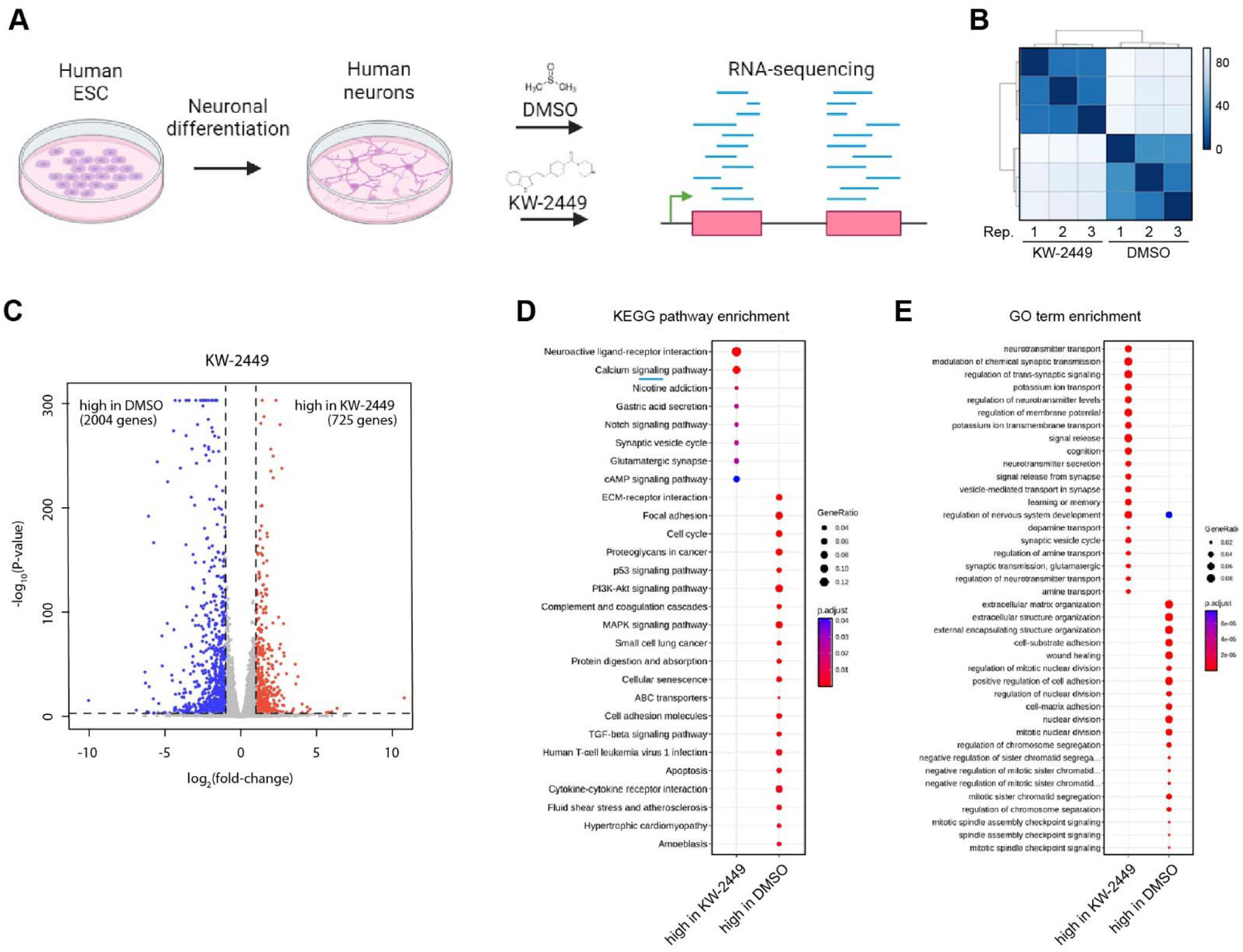
KW-2449 treatment of human stem cell-derived neurons induce gene expression changes comparable to mouse neurons. (**A**) Diagram showing the experimental workflow: (**B-C**) Correlation matrix dendrogram and volcano plots representation of RNA-seq results showing robust and consistent changes in gene expression induced by KW-2449 in human neurons. (**D**-**E**) KEGG and GO analysis of KW-2449-induced DEG highlight enrichment of synaptic genes and depletion of neuroinflammation genes.

**Figure 3:**
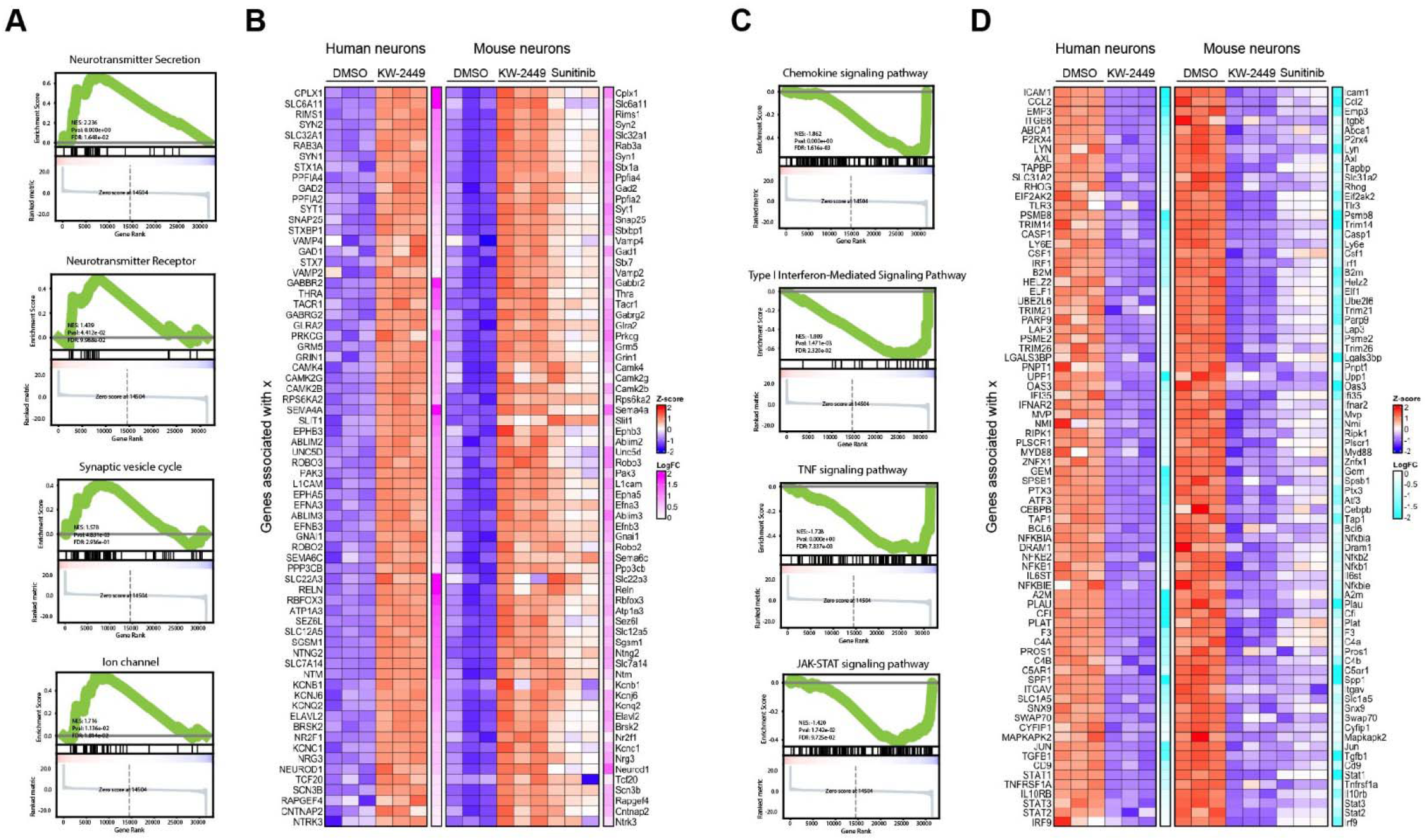
Representative gene categories regulated by FLT3 signaling in mouse and human neurons. Representative GSEA categories and curated heatmap matrix representations of the genes enriched (**A-B**) or depleted (**C-D**) in response to FLT3i drug treatment in human and mouse neurons.

### FLT3i does not alter gene expression in human microglial cells

Neuroinflammation genes are traditionally associated with immune cell types, such as microglia. FLT3 signaling in the immune system has been investigated in leukemia, but its role in the brain has not been established^*5*^. While our results show that FLT3i reduces the expression of genes associated with neuroinflammation in both mouse and human neurons, it’s possible that FLT3 signaling can also act through other cells, such as microglia, to affect neurons in a non-cell-autonomous manner. To test whether FLT3 signaling affects gene expression in human microglia, we applied KW-2449 treatment to human pluripotent stem cell-derived microglia (hMGLs) after maturation in two different microglia maturation media, referred to as Browhjohn^*22*^ and Douvaras^*23*^ protocols, and performed RNA-seq to investigate to what extent FLT3i treatment alters gene expression in these cells. In contrast to their human neuron counterparts, treating human microglia with KW-2449 does not induce significant gene expression changes, regardless of the maturation media conditions (**Fig. 4A-C**; note that, the scales of the volcano plot graphs are normalized to human neuron results shown in Fig. 2). Consistently, GSEA revealed no significant enrichment in the gene categories responsive to FLT3 inhibitor treatment in human neurons, nor was there consistent changes in the expression levels of genes responsive to FLT3 inhibitor treatment in neurons. (**Fig. 4D-G**). To understand the molecular basis of this striking difference in FLT3i treatment responsiveness between human neurons and microglia, we reasoned that microglia do not express FLT3, rendering them insensitive to FLT3i. Taken together, our results demonstrate that KW-2449 induces gene expression changes in neurons, which express the FLT3, but not in microglia which lacks FLT3. This supports the notion that KW-2449 modulates gene expression in neurons through the targeted effect of FLT3 signaling blockade.

**Figure 4:**
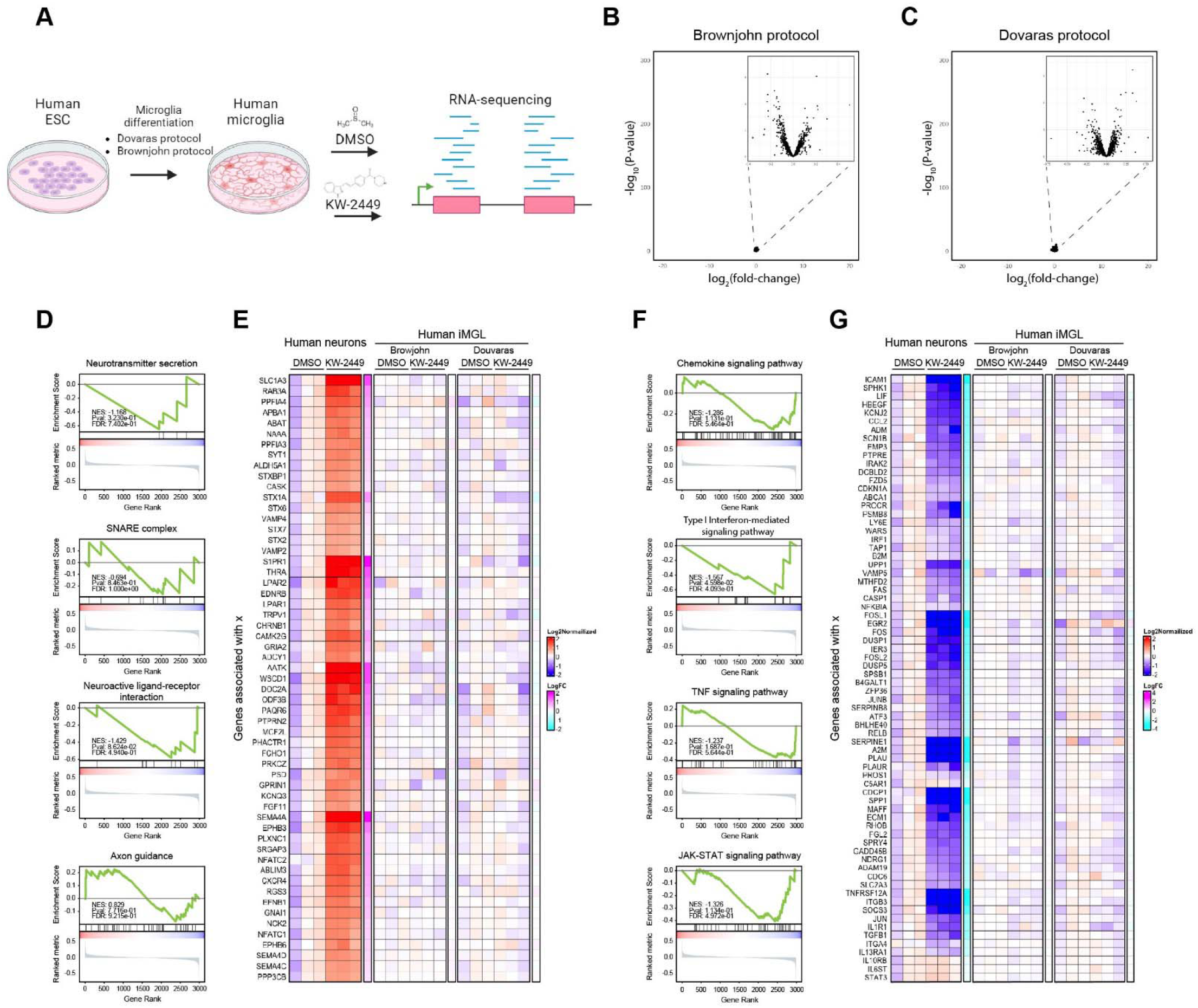
FLT3i treatment does not significantly alter gene expression in microglia. (**A-C**) Microglia generated from human pluripotent stem cells and matured with two types of maturation media do not demonstrate any significant transcriptomic change in response to KW-2449 treatment. (**D-G**) Representative GSEA plots and heatmaps comparing the genes that are expressed in both human neurons and microglia but only respond to KW-2449 treatment of human neurons but not in microglia.

### Identification of FLT3i-responsive transcription factors that regulate KCC2 expression in neurons

We further sought to elucidate the molecular pathways through which FLT3i treatment stimulates the expression of important neuronal genes such as KCC2. We hypothesized that inhibition of the FLT3 signaling pathway alters the levels of transcription factors (TFs) in neurons, one or more of which might be responsible for FLT3i-mediated upregulation of KCC2. Candidate TFs were nominated based on the following two criteria: expressed at substantial levels in neurons (defined as RPKM > 50) and, 2) undergo a more than 50% reduction in expression in response to FLT3i treatment in neurons. In order to study the function of the 18 nominated FLT3i-responsive candidate TFs at their endogenous expression levels, we experimentally knocked down these TFs individually in neurons. CRISPR/Cas9 knockdown constructs targeting the coding sequences of each candidate TF were cloned and transfected into cultured mouse cortical neurons, followed by quantitative KCC2 immunolabeling to assess their effects on KCC2 levels (**Fig. 5A**). The curated CRISPR screen revealed that knocking down several FLT3i-responsive TFs including Elf1, Elf4, Rest, Nfkb2, Stat1, Mybl1, Ezh2, and E2f8 leads to a significant increase in KCC2 expression. Therefore, FLT3i treatment-induced reduction in the expression level of these TFs may stimulate the expression of KCC2 and other genes associated with neuronal maturation. In contrast, knocking down TFs including Atf3, Scml2, Ascl1, Parp12, Irf1, Irf9 does not alter KCC2 expression, while knocking down Litaf significantly reduces KCC2 expression (**Fig. 5B-C**). Our results therefore identify a number of FLT3i-responsive TFs that may work in concert to modulate gene expression in neurons, revealing novel gene regulatory mechanisms that govern the expression of KCC2 and potentially other important genes in neurons.

**Figure 5:**
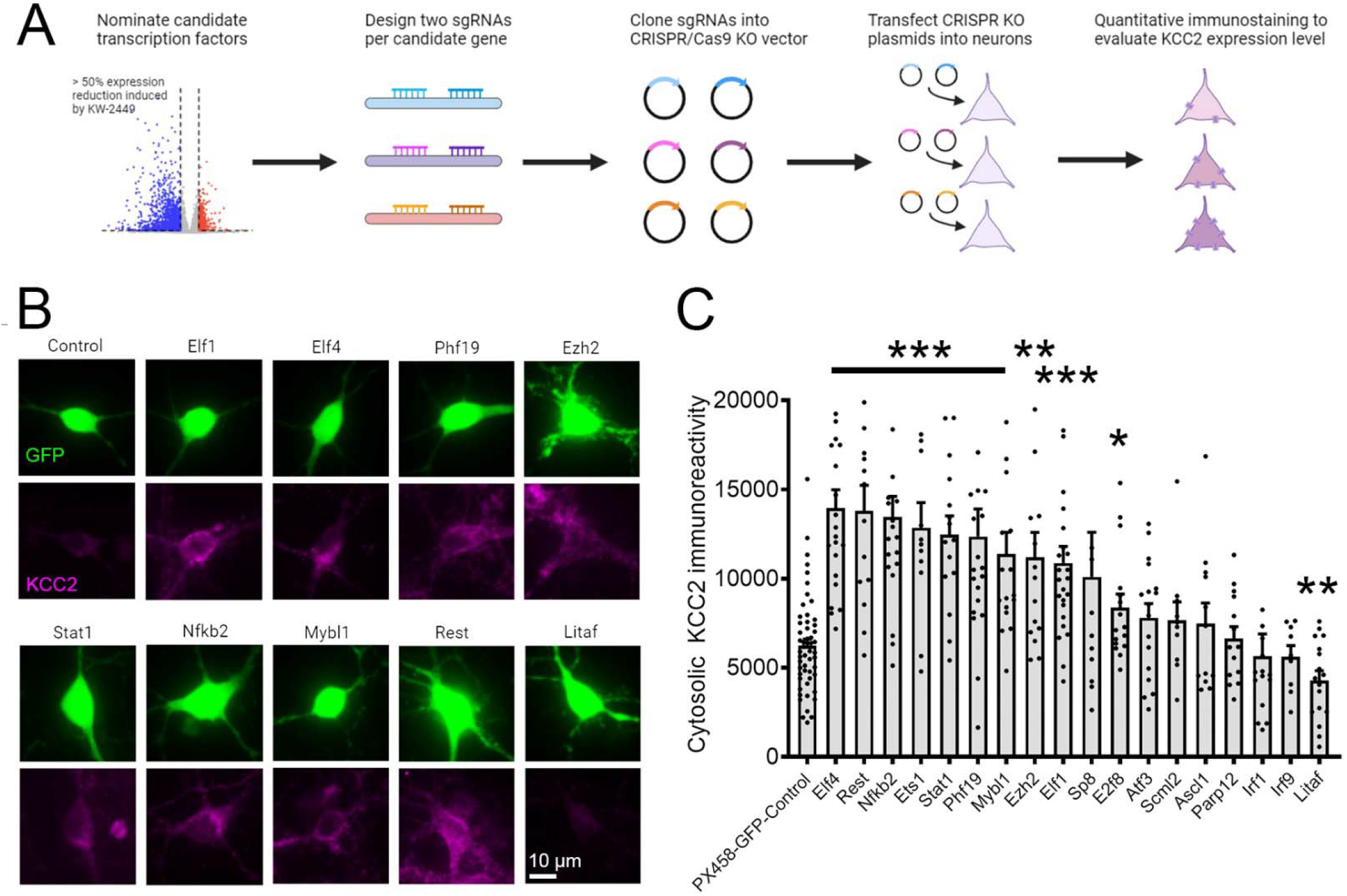
A curated CRISPR screening uncovers several transcription factors downstream of FLT3 signaling that regulate KCC2 expression. Workflow diagram showing the nomination of candidate TFs, the design and construction of CRISPR sgRNA plasmids, transfection of neurons and quantitative KCC2 immunostaining. (**B**) Representative images showing increases in KCC2 immunoreactivity in neurons that received sgRNA to knockdown candidate TFs comparing to control construct-transfected neurons. (**C**) Quantified results showing that knocking down a number of candidate TFs significantly increase KCC2 expression. Data represents mean ± SEM.

### FLT3i stimulates the expression of autism risk genes involved in the neuronal maturation processes

Cellular processes regulated by FLT3i treatment, including synaptic development and neuroinflammation, are often dysregulated in brain disorders. Mutations in genes encoding ion channels, solute transporters, and synaptic proteins are associated with increased risk for developing neurological disorders such as autism spectrum disorders (ASD). Thus, we examined to what extent FLT3i treatment changes the expression of genes in the Simons Foundation Autism Research Initiative (SFARI) autism risk gene list, and found that both KW-2449 and Sunitinib stimulate the expression of multiple ASD risk genes in both mouse and human neurons (**Fig. 6A**). We further sub-divided the genes in the SFARI list into functional categories (gene set information available upon request), and carried out GSEA analysis of each gene category’s responsiveness to FLT3i treatment. Our results show that ASD risk genes associated with synapse development, maintenance, plasticity, and ion channels, are significantly enriched in FLT3i-induced gene set, whereas genes associated with cell morphological growth, chromatin structure and transcription, and post-translational protein modification are not significantly altered (**Fig. 6B**). These results indicate that FLT3 signaling pathway may control the activities of gene regulatory networks that modulate the expression of a specific set of disease risk genes associated with neuronal functional maturation.

**Figure 6:**
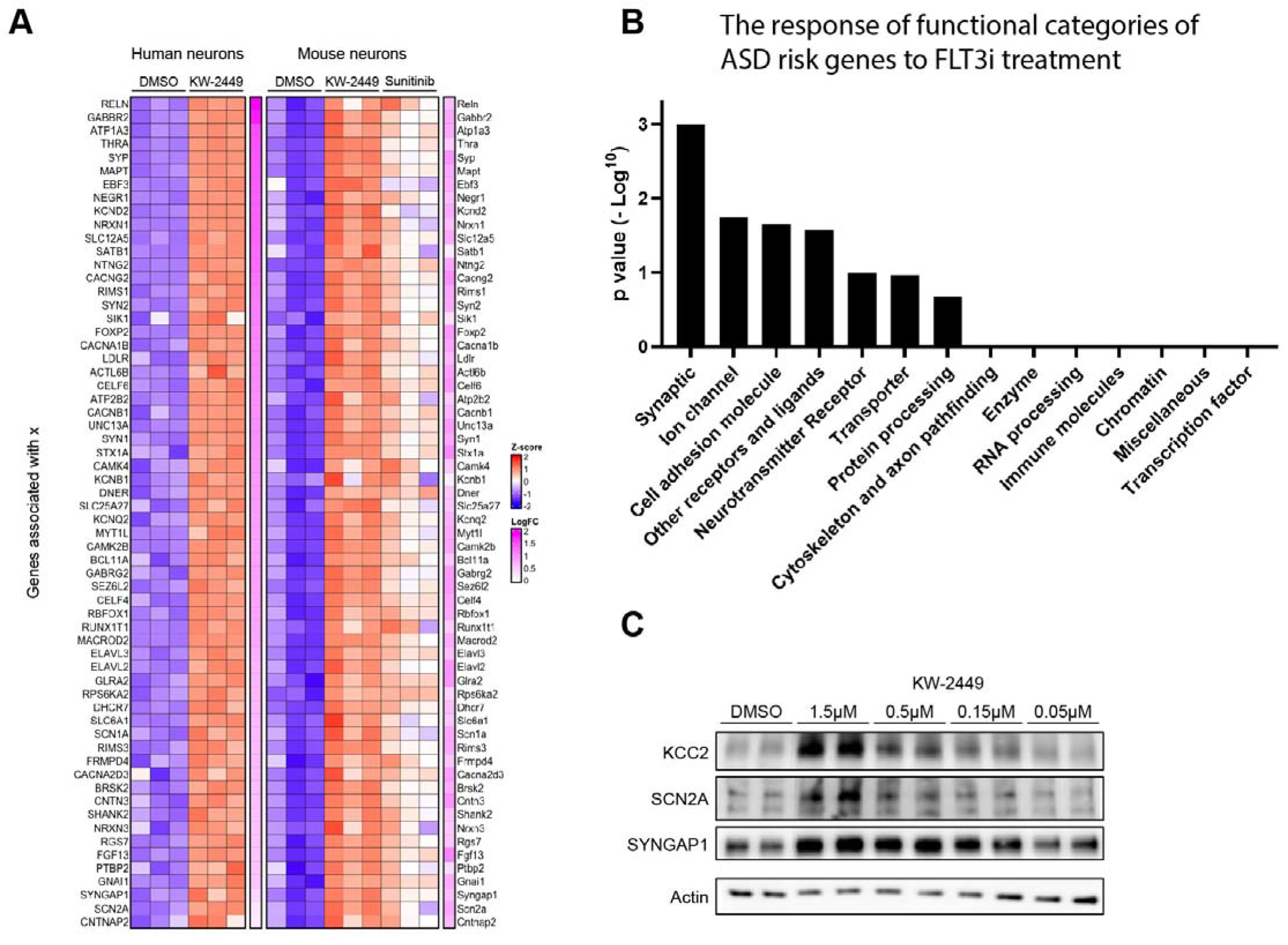
FLT3i treatment stimulates the expression of ASD risk genes. Heatmap of representative ASD risk genes stimulated by FLT3i drug treatment in human and mouse neurons. (**B**) Gene set enrichment score for the various functional categories of genes. (**C**) Validation of FLT3i-induced stimulation of ASD risk gene expression at the protein level with Western blot method.

Representative FLT3i-induced disease risk genes encode functionally important proteins including the ion channel proteins SCN1A and SCN2A, solute transporters KCC2 (SLC12A5) and GABA transporter (SLC6A1), as well as synaptic proteins SYNGAP, SHANK2, and NRX. Insufficient level of NDD risk gene products in the brain, due to genetic mutations that cause haploinsufficiency or epigenetic silencing that suppresses gene expression, is a common mechanism of NDD pathogenesis, and herefore, FLT3 may be leveraged as a target for therapeutic development. Thus, we examined the validity of developing FLT3i as a therapeutic approach for restoring NDD risk gene expression through biochemical validation of a number of FLT3i-responsive NDD risk genes, including ion channel SCN2A, chloride transporter KCC2, and synaptic protein SYNGAP. Our results show that treating mouse primary brain cell culture with the FLT3i drug KW-2449 significantly enhances the protein levels of KCC2, SYNGAP, and SCN2A in a dose-dependent manner (**Fig. 6C**). These results validate our RNA sequencing results at the protein level, and provide the rationale for using FLT3i drugs to rescue deficiencies in NDD-associated genes.

### KW-2449 administered in mouse model can enter the brain to enhance KCC2 protein expression

Encouraged by the *in vitro* results demonstrating the efficacy of KW-2449 in stimulating the expression of KCC2 and other genes critical for brain health, we further examined the brain penetrance and *in vivo* target engagement of the FLT3i drug KW-2449 in mice. Our previous work demonstrated that application of KW-2449 rescues behavioral phenotypes caused by reduced KCC2 expression in a mouse models of Rett syndrome^*17*^. Pharmacokinetics/pharmacodynamics (PK/PD) experiments were conducted to administer KW-2449 through either intraperitoneal (IP) or oral delivery (PO) routes. Blood or brain samples were collected at various time points following KW-2449 administration, and liquid chromatography (LC) analysis were conducted to determine the concentration of KW-2449 in biological samples (**Fig. 7A**). Our results show that KW-2449 delivered to mice through either the IP or PO route can enter the bloodstream and is metabolized with a half-life of approximately three hours (**Fig. 7B**). Our previous work showed that KW-2449 treatment of cultured human neurons or mouse brain slices elevates KCC2 expression and restores GABAergic inhibition in a model of Rett syndrome^*24*^. Importantly, our results show that KW-2449 readily enters brain tissue at a concentration of 220 ng/ml, which is within the effective dose range found to stimulate KCC2 expression in neurons *in vitro* (**Fig. 7B**)^*17*^. In the brains of mice receiving KW-2449 injections, we detected a substantial elevation in KCC2 protein expression in cortex. Intriguingly, a single IP administration of KW-2449 stimulates KCC2 expression starting at six hours and persisting for up to three days (**Fig. 7C**). Such a long-lasting effect of KW-2449-induced KCC2 expression even after the drug is mostly cleared from the bloodstream suggests sustained activation of a signaling cascade induced by drug treatment leading to activated *KCC2* gene expression in neurons, combined with the long half-life of KCC2 protein in neurons, which indicates that the drug can be applied at long intervals and still achieve therapeutic benefit. Thus, our results show that FLT3i compound KW-2449 can cross the blood-brain barrier to enter the brain and engage the molecular target to enhance KCC2 expression for a prolonged period of time.

**Fig. 7:**
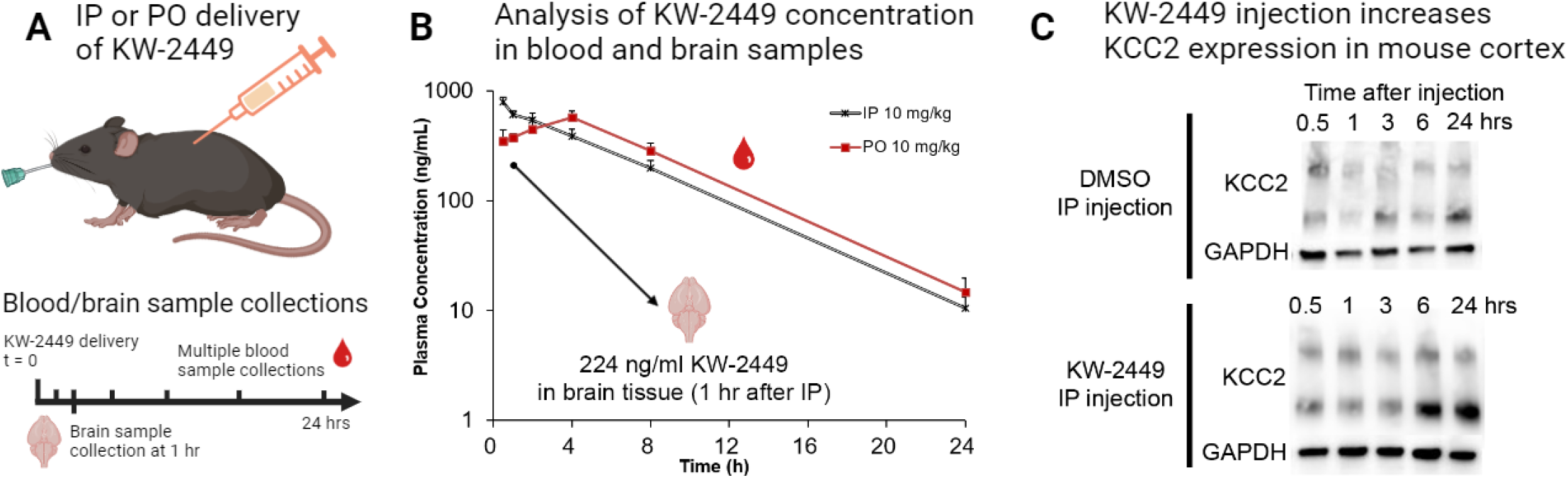
KW-2449 stimulates KCC2 expression *in vivo*. (**A-B**) Experimental diagram and results of PK/PD studies show that KW-2449 is metabolized in the blood with a half-life of ∼ 3 hours, and can enter the brain at an effective concentration. (**C**) I.P. delivery of KW-2449 induces KCC2 expression in the mouse cortex six hours after injection and lasts for 24 hrs.

### KW-2449 treatment suppresses kainic acid-induced EEG seizure *in vivo*

KCC2 deficiency is a key molecular event leading to the onset and recurrence of drug-resistant epilepsy^*25*^. We hypothesize that the increase in KCC2 expression induced by KW-2449 treatment will strengthen GABAergic inhibition and exert anti-seizure effect *in vivo*. To test this hypothesis, we conducted electroencephalography (EEG) recording experiments in a chemoconvulsants kainic acid (KA)-induced model of pharmacoresistant epilepsy. Wild-type mice were implanted with EEG recording electrodes and randomly assigned to two groups and received injections with either 2 mg/kg KW-2449 or DMSO vehicle control. We then performed KA challenge and EEG recording 24 hours after KW-2449 injection, a time point when the KCC2 expression in the brain is significantly enhanced (**Fig. 8A**). Compared to DMSO-injected control mice, pre-treatment of KA-induced mice with KW-2449 significantly reduces the duration of seizures, delays the onset of seizure activity and status epilepticus (SE), which are prolonged epileptic burst activities damaging to the brain circuits^*26*^, (**Fig. 8B-C**). Importantly, SE can be prevented in more than 70% of experimental mice injected with KW-2449, while all the mice injected with DMSO vehicle develop SE after KA challenge (**Fig. 8C**). Taken together, our results demonstrate that FLT3i drug KW-2449 can enter the brain to increase KCC2 expression and reduce seizure susceptibility in a mouse model of refractory epilepsy.

**Fig. 8:**
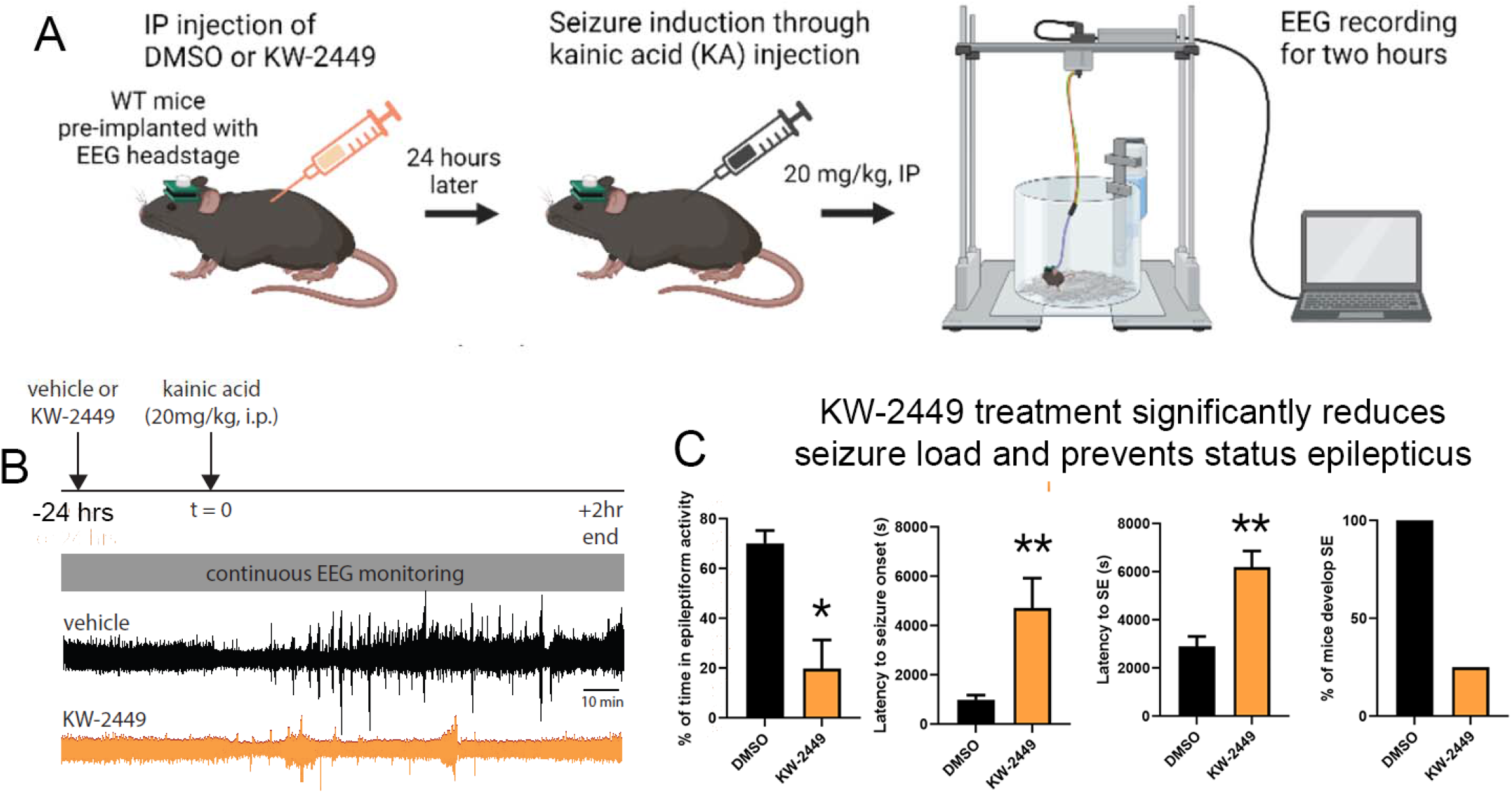
KW-2449 stimulates KCC2 expression *in vivo* and reduces seizure susceptibility. Experimental diagram, representative EEG recording, and quantified results show that KW-2449 treatment of mice 24 hours prior to the KA challenge significantly delays seizure onset and reduces seizure load. (^*^ p < 0.05, ^**^ p < 0.01. n = 5 mice for vehicle and 7 mice for KW-2449-injected groups). In all panels, data represents mean ± SEM.

## Discussion

In this study, we utilized a pharmacogenomics approach to elucidate the effects of FLT3 kinase inhibitors, previously identified in our drug screen as beneficial for rescuing Rett syndrome phenotypes, on both murine and human brain cells as well as in a mouse model of epilepsy. Our results highlight previously unknown roles of FLT3 kinase signaling in regulating the expression of genes important for brain development and neuroinflammation in neurons, but not in microglia that do not express the FLT3 receptor. Traditionally, FLT3 has been studied in blood cell development and FLT3i drugs were developed for the treatment of leukemia^*5*^.

Expression of FLT3 mRNA and protein in the brain has been reported^*8*^, and several recent studies point to a possible role of FLT3 in regulating neuronal function^*15*^; however, its function in the brain has not been previously investigated, and to our knowledge, this study provides the first systematic demonstration of the functional significance of FLT3 signaling in the developing and mature brain.

Our results show that the FLT3i treatment regulates a similar set of biological processes in both cultured human and mouse neurons, including synaptic development, neuronal morphological and functional maturation, and neuroinflammation. Such cross-species consistency indicates that the key molecular mechanisms driven by FLT3 signaling in the brain are evolutionarily conserved and functionally meaningful. While the general pathways are conserved, specific genes may be differentially regulated by FLT3i in human and mouse neurons. Considering that the human neurons used for this study were differentiated from stem cells whereas the mouse neurons were cultured from neonatal mouse brain, the distinctions in FLT3i responsiveness could be due to differences in cell composition in the culture, the maturation level of neurons, or species-specific features. To begin to address this difference and dissect the cell type-specific response to FLT3i, we demonstrated that KW-2449 treatment does not induce any gene expression changes in human stem cell-derived microglia. We further show that microglia do not express FLT3 kinase while the neurons do. The neuron-specific responsiveness to KW-2449 indicates that the effect that KW-2449 exerts on neurons is likely to be mediated by an on-target mechanism of FLT3i.

Leveraging the RNA-seq data, we nominated candidate TFs downstream of FLT3 signaling that may regulate gene expression in neurons, and conducted a curated CRISPR screening study to knockdown their endogenous expression levels. This work has led to the identification of several TFs that respond to FLT3i and that regulate KCC2 expression levels in neurons under physiological conditions. One of the identified TFs is Signal transducer and activator of transcription 1 (Stat1), a TF responsive to interferon-gamma (IFN-γ) signaling to mediate pro-inflammatory functions in various non-brain cell types. In the context of the brain, Stat1 activation in neurons after ischemia exacerbates brain injury^*27*^, while *Stat1* knockout mice show improved spatial memory formation^*28*^. Our findings elucidate a molecular mechanism by which Stat1 activity is modulated in neurons, subsequently influencing neuronal function. Two other identified TFs, enhancer of zeste homolog 2 (Ezh2) and PHD finger protein 19 (Phf19), are components of the Polycomb Repressor Complex-2 (PRC2).

Our discovery of an epigenetic mechanism wherein FLT3i downregulates the expression of PRC2 complex components Ezh2 and Phf19, which presumably lead to reduced deposition of H3K27me3 repressive marks on target genes, aligns with findings from a previous study conducted on leukemia cells^*29*^. In the context of brain development, a recent study reports that EZH2 is a master suppressor of the expression of genes associated with neuronal functional maturation in human neurons^*30*^. Another study in mice discovered that Ezh2 expression levels are upregulated following status epilepticus and that pharmacological inhibition of Ezh2 increases seizure burden^*31*^. The seemingly opposite effect of FLT3i and Ezh2i drugs in regulating seizure propensity may be due to the differences in the cell types responding to drug treatment: Unlike EZH2, for which expression is low in the brain but high in many non-neuronal cell types, FLT3 expression is enriched in the brain, especially in neurons, relative to most tissues and cell types (Human Protein Atlas). Moreover, *Ezh2* KO leads to lethality at early stages of mouse development^*32*^. Therefore, indirect modulation of PRC2 complex activity in the brain through FLT3i may present a safer and more effective therapeutic approach. RE1 silencing transcription factor (REST) has a known function in suppressing KCC2 expression in neurons^*33*^, a finding that is reproduced in our study. Other TFs identified from the screening, including Elf1, Elf4, Nfkb2, Myb1l, and E2f8, have not been studied extensively in the brain. Our results uncover previously unknown functions of these TFs in regulating important neuronal gene such as KCC2, which warrants further investigation of their roles in brain development and functioning. A limitation of our current analysis is that the candidate transcription factors (TFs) were nominated based on their changes in expression levels in response to FLT3i treatment, potentially overlooking TFs that undergo phosphorylation or activity level changes following FLT3i exposure. Future studies using complementary approaches such as phosphoproteomics could provide a more comprehensive mechanistic understanding of the scope of FLT3 signaling in neurons.

Unlike most current anti-seizure medications that require direct binding of the drug to the target protein, FLT3 inhibitors modulate the mRNA and protein expression of disease-relevant genes. This mechanism allows these drugs to target traditionally non-druggable proteins, offering possibility of pharmacologically enhancing the expression and activity of ion channels, transporters, and synaptic proteins. One important FLT3 target gene is neuronal chloride transporter KCC2 that plays a critical role in maintaining GABAergic inhibition in the brain.

KCC2 down-regulation facilitates epileptic seizures^*34*^, while knocking out KCC2 induces severe motor deficits in mice^*35*^; conversely, viral overexpression of KCC2 in hippocampal pyramidal cells reinstates inhibition and reduces seizures in a chemoconvulsant-induced mouse seizure model^*36*^. We performed *in vivo* pharmacokinetics and pharmacodynamics (PK/PD) studies to characterize the brain availability of KW-2449, as well as target engagement experiments to demonstrate the KW-2449 induces KCC2 expression in the mouse brain. To test the anti-seizure efficacy of KW-2449, we used the KA-induced seizures in mice for our study because it provides a clinical relevant model of limbic/generalized epilepsy and hippocampal damage caused by domoic acid exposure in human patients^*37*^; also, the KA-induced model recapitulates disease-relevant abnormalities in electrophysiology, neurochemistry, and brain structure commonly observed in human pharmaco-resistant seizure patients. Results from *in vivo* pharmacology and EEG recording experiments show that KW-2449 treatment reduces seizure burden and suppresses status epilepticus in the KA-induced mouse model of pharmacoresistant epilepsy. These promising *in vivo* results highlight the potential of FLT3 inhibitor KW-2449 to be further developed and translated as a novel disease-modifying agent for treating patients who do not respond to any of the currently available anti-seizure medications.

Our work identified a set of disease risk genes associated with autism spectrum disorder and epilepsy, for which expression levels are upregulated by FLT3i in neurons. This result raises the possibility that FLT3i drugs may potentially serve as broadly applicable therapeutics in cases of NDD-associated gene haploinsufficiency, where half of the protein expression is lost due to *de novo* heterozygous mutations, through stimulating gene expression output from the intact wild-type allele. The spectrum of clinically-identified mutations in NDD-associated genes, including in *SYNGAP1* and *SCN2A*, comprises mainly monoallelic nonsense mutations that truncate and inactivate their protein products^*38*^. Preclinical studies show that re-expression of the Syngap1 protein through mouse genetics in a model of *Syngap1* haploinsufficiency^*39*^, or reactivation of *Scn2a* gene expression in adult *Scn2a*^*+/-*^ mice through gene therapy^*40*^, improves brain function and rescues seizure and cognitive impairments. These findings indicate that restoration of SYNGAP or SCN2A expression through FLT3i drug treatment may lead to functional recovery even after the onset of disease symptoms in patients.

Our profiling of FLT3i-responsive disease risk genes lays the groundwork for stratifying and pre-selecting patients based on their genotypes, potentially identifying those who may benefit most from FLT3i drug treatment.

A feature of FLT3i drug is that it modulates the expression of multiple genes in neurons, presumably by modulating a gene co-regulatory network through various FLT3i-responsive transcription factors. The safety and off-target effect of such gene-modulation drugs need to be carefully examined. Kinase inhibitor drugs that modulate expression of multiple proteins, such as Evolimus that inhibit the mTOR pathway, have been developed for treating refractory seizures^*41*^. KW-2449 has been tested in clinical trials for treating leukemia and has shown good safety profiles in patients. Our results show that a single administration of KW-2449 at a dose substantially lower than that used in cancer therapy elevates brain KCC2 expression for at least 24 hours.

Such a prolonged time course is favorable for clinical application because it is not necessary to maintain a constant high drug concentration. Therefore, a sparse dosing regimen can be applied to reduce toxicity and adverse effects. To establish the safety and feasibility of a novel therapeutic approach that targets FLT3 signaling to normalize expression of NDD-associated genes and rescue disease phenotypes, further long-term toxicology studies are required to assess the effects of FLT3i drugs on brain and non-brain tissue, and to test FLT3i in various genetic mouse models of epilepsy and neurodevelopmental disorders. Future studies may focus on analyzing large-scale human genetic and functional genomics datasets to investigate to what extent FLT3 loss-of-function variants exert disease-modifying effects that suppress disease symptoms in patients harboring mutations in NDD-associated genes.

In Summary, our work provides the first demonstration the functional significance of FLT3 kinase in brain development and in maturation, revealing previously unknown cellular and molecular mechanisms of FLT3 signaling in brain cells, and profiles NDD risk genes responsive to FLT3i drug treatment in neurons. We further demonstrate that the small molecule FLT3i compound KW-2449 is orally bioavailable, readily enters the brain to increase KCC2 expression levels, and exerts an anti-seizure effect in two models of refractory epilepsy. Our results warrant further mechanistic investigation to elucidate the role of the FLT3 signaling cascade at different stages of brain development, and to establish the basis for the development of novel anti-epileptic drugs that correct KCC2 dysregulation to restore GABAergic inhibition in an epileptic brain.

## Methods

### Mouse and human neuron culture

To prepare primary mouse cortical neuron culture, neonatal mouse brains were extracted from BL6/C57 mice at postnatal day 0-2 through dissection, and the cerebral cortex tissue were isolated, cut into small pieces, and digested with papain for five minutes. Trypsin inhibitor was added to stop the digestion reaction, and the tissue pieces were then rinsed twice and underwent mechanical dissociation with a P1000 pipette. The dissociated cell suspension was then filtered through a cell strainer to remove small tissue clumps, and seeded at a density of approximately 100,000 cells per well into matrigel-coated 12-well plates for sequencing and biochemistry assays, or at approximately 50,000 cells per well onto 12 mm diameter glass coverslips coated with poly-D-lysine followed by Matrigel placed in 24-well plates for transfection and imaging assays.

To generate human pluripotent stem cell-derived neuron culture, we adapt a previously published protocol^*17*^. Briefly, neural progenitor cells (NPC) were generated from *MECP2* knockout WIBR3 human embryonic stem cells^*42*^. NPC were seeded at a density of approximately five million cells per flask onto T75 flasks and undergo neuronal differentiation and maturation for six weeks. The neurons were then re-plated into 6-well plate at a density of about 500,000 cells per well, matured for another two weeks before drug treatment and RNA collection. For immunostaining and imaging analysis, NPCs were seeded at 10,000 cells per well into 12 mm coverslips pre-seeded a mouse astrocyte feeder layer to support maturation^*43*^. The neuronal cultures were then kept in a neuronal maturation medium for six to eight weeks before analysis.

### Human microglia culture and co-culture with neurons

Human pluripotent stem cell-derived microglia-like cells (hMGLs) were generated with H1 embryonic stem cells^*44*^, using two published protocols^*22, 23*^ with the following modifications: for the Brownjohn protocol, embryoid bodies were plated into 10 cm matrigel-coated dishes, and floating primitive macrophage precursors (PMPs) were later collected and plated in 6 well tissue culture-treated plates in one of two maturation media: NGD base media^*44*^ supplemented with 50ng/mL TGF-β, 100ng/mL IL-34 and 25ng/mL M-CSF, or Neurobasal-based media with 1X GlutaMAX, 1X Gem21 without vitamin A, 50ng/mL TGF-β, 100ng/mL IL-34 and 100ng/mL M-CSF. After 7 days in maturation media, cells were harvested for RNA extraction. For the Douvaras et al protocol, ∼100-200 colonies were plated in matrigel-coated 10cm dishes in mTeSR Plus, and STEP1 media was added once the average colony size was ∼1mm in diameter.

### FLT3i drug treatment

FLT3i drugs were dissolved in DMSO and applied to the culture medium of neurons or microglia cells at a final concentration of 2 µM for KW-2449, and 1 µM for Sunitinib. The same volume of DMSO were added to parallel cultures as control. Three days after drug treatment, total RNA samples were prepared for sequencing. For the *in vivo* administration of KW-2449, 10 mg of KW 2449 (MedChem Express, HY-10339) was dissolved in 100 µl DMSO to make 100 mg/ml stock solution, which were then aliquot into sterile Eppendorf tubes at 10 µl/tube and stored at -20°C. Male BL6/C57 mice were purchased from the Jackson Lab. On the day of the drug administration, 2 µl of the KW-2449 stock solution is thawed and diluted into 1 ml sterile saline at room temperature. Vortex until the initial white precipitate are fully dissolved. The same amount of DMSO was dissolved in 1 ml saline as control. Mice were IP injected with 10 µl/g body weight with KW-2449 or DMSO working solution. The final concentration of KW 2449 used for IP injection is 2 mg/kg bodyweight in 0.2% DMSO.

### RNA sequencing and data analysis

Total RNA was extracted from cultured neurons or microglia with the Qiagen RNeasy® Micro kit and then submitted to the Whitehead genomics core facility and the Azenta for cDNA library construction and sequencing using standard next generation sequencing (NGS) protocols. The NGS sequencing results were then mapped to human or mouse genome. Count matrices (genes by sample) were prepared using nf-co.re/rnaseq v.3.10.11. The hg38 genome was used as a reference for human samples, while the mm10 genome was used for mouse samples. Differential expression analysis was performed using DESeq2^*45*^. A gene was considered differentially expressed if its absolute log2 fold-change was greater than 1 and its adjusted p-value was less than 0.5. To identify enriched gene sets, gene set enrichment analysis (GSEA) was conducted using the KEGG pathways and GO:BP databases as references^*46, 47*^. The fgsea R package was used, and the computed fold-changes were used as the metric value5. In addition, enrichment analysis based on the hypergeometric test was performed using ClusterProfiler6. A gene set was considered significant if its adjusted p-value was below 0.05. The functionally annotated ASD risk gene lists were curated based on the functional categories of individual SFARI risk genes available upon request. We used a GSEApy use a permutation test to calculate p-value/FDR and sometimes it can be zero. They suggest using 1/(number of permutations) number to replace the 0 values (https://gseapy.readthedocs.io/en/latest/faq.html#q-why-p-value-or-fdr-is-0-not-a-very-small-number). We are using default number of permutations which is 1000, so we can replace 0 values with 0.001.

### Candidate transcription factor nomination and CRISPR screening

The candidate transcription factor that may mediate the FLT3i treatment stimulates the expression of important neuronal genes such as KCC2, we performed bioinformatic analysis to nominate candidate transcription factor (TF) that may drive such transcriptional program based on two criteria: 1) TFs that are expressed at substantial levels in neurons (defined as RPKM > 50 in mouse neurons) and 2) underwent > 50% reduction in average gene expression in response to FLT3i treatment. Two knockout sgRNAs sequences targeting the obligatory coding exon were selected per candidate TF gene from the mouse GecKO library^*48*^ and cloned into a PX458 SpCas9-GFP CRISPR knockout construct^*49*^.

The candidate TF knockdown plasmids were then transfected into cultured mouse neurons at 3 days-in-vitro (DIV 3) to knockout candidate TF in respective biological samples. Plasmids are mixed with Lipofectamine 2000 reagent at a 1 µg/2.2 µl ratio for each coverslip. PX458 SpCas9-GFP backbone construct was included as a non-targeting negative control in each batch of the experiment. One week after the transfection, KCC2 immunoreactivity in the cytosol and major dendrites of GFP positive transfected neurons are quantified as a readout for the screening. We choose the time point of DIV 10 to assess the KCC2 expression changes in mouse neurons because at this developmental time point, KCC2 protein is expressed but have not reached high level as they are in mature neurons, enabling the detection of both positive and negative regulators of KCC2 gene expression.

### Immunocytochemistry

Co-cultured human iPSC neurons and human iPSC microglial cells with tdTomato labeling were fixed on coverslips with 4% PFA for 15 minutes at room temperature (RT) and washed with PBS. Cell coverslips were washed in PBS once and 10% Normal Donkey Serum (NDS) and 2% Bovine Serum Albumin (BSA) in PBS-T (0.2% Triton X100) were applied for 30 minutes to block non-specific binding. The cells were then incubated with c-terminal rabbit anti-Flt3 primary antibody (1:200 dilution), chicken anti-mCherry (1:200 dilution, to boost tdTomato signals) and mouse anti-beta III tubulin (TUJ1, a neuronal marker, 1:500 dilution) primary antibodies diluted in blocking buffer at 4°C overnight. The negative control used the same combination of primary antibodies with the exception of rabbit IgG replacing the Flt3 antibody. After washing cells 4 times with PBS the next day, the cells were incubated with for 2 hours RT with secondary antibodies including donkey anti-rabbit 647, donkey anti-chicken 594 and donkey anti-mouse 488 secondary antibodies, diluted 1:500 in blocking buffer. Afterwards the coverslips were washed 3 times with PBS for 5 minutes each and mounted with DAPI-fluoromount mounting medium.

### Image analysis

Images were analyzed using ImageJ/Fiji software. Images were opened in ImageJ and using the ROI Manager, regions of interests (ROIs) were drawn around the cell body and a portion of a major dendrite using the GFP channel of the GFP-positive (successfully transfected) cell. Next, using the DAPI channel, an ROI around the nucleus of the GFP positive cell was drawn. Using these three ROIs, the area and average intensity of each was measured through the 647 channel which showed the KCC2 staining. The background intensity of the KCC2 image was recorded as well. To compute the KCC2 intensity on the cellular membrane (*I*_*cm*_), the equation below was used:

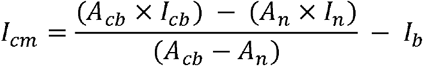

where *A*_*cb*_ and *I*_*cb*_ are the recorded area and average KCC2 intensity of the cell body respectively, *A*_>*n*_ *I*_>*n*_are the area and average KCC2 intensity of the nucleus respectively, and *I*_>*b*_ is the background intensity. To compute the intensity of KCC2 on the major dendrites (*I*_>*d*_), the equation below was used:

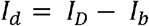

where *I*_D_ is the measured KCC2 intensity in the major dendrite and *I*_b_ is the measured background intensity. The area of the dendrite ROI is not needed for calculations.

### PK/PD measurements

The PK/PD experiments were performed with the service of a CRO company (HD Biosciences). Male CD1 mice weighing 28-30 grams were used for the study. Two routes of administration (PO and IP) were tested, and blood samples were collected at six time points: 0.5, 1, 2, 4, 8, 24 hours post administration. Three mice were evaluated in each administration route/time point group. Mice in the PO groups have *ad lib* access to water and were fasted overnight before dosing and free access to food at 4hr post dosing. Mice in the IP groups have free access to food and water. KW-2449 was formulated at 10 mg/kg in 5% DMSO + 10% Solutol^®^ HS 15 (solubility enhancer) + 85% water. The volume of both IP and PO administration is 10 ml/kg. Three mice were dosed with 10 mg/kg KW-2449 through the IP route and their brain sample were collected at 1 hr post injection. For plasma and brain sample collection, the animals were restrained manually at the designated time points, approximately 110 µL of blood sample was collected via facial vein bleeding into K2EDTA tubes. The blood samples were maintained in wet ice first and centrifuged to obtain plasma (2000g, 4_°C_, 5 min) within 15 minutes post sampling. The brains from the animals were excised, brain samples were collected and weighted, and then snap frozen on dry ice and further stored at -70 °C.

An UPLC (Waters) chromatographic system equipped with an API6500 QTRAP mass spectrometer (Applied Biosystems, Concord, Ontario, Canada), and an Analyst 1.6 software packages (Applied Biosystems), were used to control the LC-MS/MS system for data acquisition and processing. The DMSO stock solution was diluted with 70% acetonitrile to get working solutions of STDs and QCs, then mixed 3 µL working solution with 27 µL blank plasma to get STD and QC samples. An aliquot of 30 μL plasma was spiked into a 1.5 mL tube, and 150 μL of acetonitrile containing internal standard were added for protein precipitation. The mixture was vortexed, centrifuged at 14000 rpm for 5 min. Transfer 100 μL supernatant into 96 well plate, and 300 μL water was added, the mixture was vortexed and injected for LC-MS/MS analysis. Brain sample was homogenized with ice-cold phosphate buffer saline (pH 7.4) at a ratio of 4 (buffer) : 1 (tissue) (v/w). An aliquot of 30 μL homogenate was spiked into 1.5 mL tube, and 150 μL of acetonitrile containing internal standard was added for protein precipitation. The mixture was vortexed and centrifuged at 14000 rpm for 10 min. 100 μL of supernatant was mixed with 300 μL of water and the final solution was injected for LC-MS/MS analysis. All data are acquired using Analyst 1.7 software (Applied Biosystems). Plasma concentration versus time data will be analyzed by non-compartmental approaches using the WinNonlin software program (version 6.1, Pharsight, Mountain View, CA).

### Western blot

Primary mouse neurons were cultured for 3 days before KW-2449 drug treatment, which were administered for 6 days, and neurons were collected with Pierce Lysis Buffer on DIV9. Samples were sonicated briefly for 5 seconds and centrifuged at 16,000g for 10min at 4C. Supernatant was collected and mixed with BioRad Laemmli sample buffer at a 3:1 ratio, then heated for 5min at 95C. Samples were run on western blot gels from BioRad at 300V for approximately 20min. Gels were transferred using the TurboBlot system from BioRad. Membranes were then blotted with primary antibody at 4C overnight, then secondary antibody for 1 hr at room temperature. Membranes were then visualized using an Azure chemiluminescent imager with Clarity ECL substrate (BioRad).

The mouse brain samples were collected at different time points after KW-2449 injection. Adult C57 mice (8-12 weeks old, male) received either compound KW2449 (2 µg/g bodyweight in 0.2% DMSO/saline) or 0.2% DMSO/saline, IP. Mice from each treatment condition were randomly picked up, according to the different time points, to be perfused with 4°C cold saline to wash off blood, and their brain motor cortex samples were immediately harvested and saved in dry ice, then transferred to -20°C freezer. The brain samples were lysed with RIPA buffer with cOmplete protease inhibitor cocktail tablet (1 tablet/10 ml buffer), sonicated, and centrifuged for 5 min, 3000 rpm at 4°C, then the supernatants were used for western blots. The proteins were separated in 4-20% gradient gels (BioRad) and transferred in nitrocellulose membranes. After blocking with EveryBlot blocking buffer (Bio-Rad, Cat#: 12010020), the membranes were incubated with anti-KCC2 (rabbit polyclonal, 1:1000, Sigma, Cat#: 07-432) or anti-GAPDH (mouse, 1:1000, abcam, Cat#: 125247) respectively, in blocking buffer, 4°C overnight. After TBS-T washing, the membranes were incubated with Donkey anti-rabbit-or anti-mouse HRP-conjugated secondary antibodies (1:500) for 2 hours at room temperature. ECL (Bio-Rad, Cat#:1705061) was used as the substrate and Azure Imaging system was used for imaging

### Electroencephalogram (EEG) recording

Seizure susceptibility in mice was evaluated based on electrographic seizure activity measured using electroencephalogram (EEG) recording. Mice were implanted with chronic EEG headmounts under anesthesia with 100 mg/kg ketamine and 10 mg/kg xylazine. A lengthwise incision was made along the scalp and a pre-fabricated headmount (Pinnacle Technology, part #8201) was fixed to the skull with four screws, two of which serve as EEG leads, one a reference ground, and one an animal ground. The site was closed with dental cement and the mice were allowed to recover for a minimum of a week prior to experimentation.

Electroencephalogram recordings were collected using a 100x gain preamplifier high pass filtered at 1.0 Hz (Pinnacle Technology, part #8202-SE) and tethered turnkey system (Pinnacle Technology, part #8200). Data acquisition was performed using a Powerlab system (ADInstruments) and LabChart Pro software (ADInstruments). Mice were treated with KW-2449 (2mg/kg, i.p.) either 30 mins or 24 hours prior to kainic acid administration. Electrographic activity was recorded for two hours following administration of kainic acid (KA, 20mg/kg, i.p.). The latency to the first ictal event and total time exhibiting abnormal electrographic epileptiform activity during the two-hour recording period following kainic acid administration and one hour following diazepam treatments was measured. Abnormal epileptiform activity was detected as paroxysmal activity having a sudden onset and an amplitude at least 2.5x the standard deviation of the baseline and a consistent change in the Power of the fast Fourier transform of the EEG, including a change in the Power and the frequency of activity over the course of the event. These parameters have been used previously by our laboratory and others to measure electrographic epileptiform activity. Abnormal electrographic activity, including ictal activity and rhythmic spiking lasting longer than 30 s were collectively defined as “epileptiform activity”. The cumulative amount of time exhibiting epileptiform activity was divided by the length of the total recording period × 100 was calculated as the “% time epileptiform activity”. Kainic acid is an established model of temporal lobe epilepsy in rodents in which seizure activity progresses to status epilepticus (SE) and development of pharmacoresistance^*26*^. In this model, animals enter SE, defined as unremitting, persistent epileptiform activity lasting at least 5 mins. The latency to SE is defined as the latency from kainic acid administration to the start of SE. The percentage of animals that progress to SE and the percent mortality was also documented.

### Statistics

We are committed to full transparency, i.e. reporting our findings regardless of the experimental outcome. The following steps will be employed to produce robust experimental results: Individual cells were selected for imaging or recording analysis regardless of the genotype/treatment condition based on cell morphology that does not reveal key electrophysiological parameters. When performing *in vivo* measurements such as video/EEG analysis of seizure phenotypes and rescue thereof by KW-2449 treatment, the researchers were blinded as to the drug treatment conditions when collecting and analyzing data. Data sets were tested for normality and equal variance and, if those criteria are met, Student’s t-test was used. If the criteria are not met, a non-parametric Mann-Whitney test was used. For multiple group comparisons one-way ANOVA with Bonferroni post hoc test were used. Differences were considered statistically significant at p < 0.05. Data were compiled from at least three experiments and reported as mean ± SEM.

## Author contributions

X.T., J.M., and M.G. designed research; K.C., M.M., M.G., E.C., Y.Y., S.L., E.S., T.L., and X.T. performed research; S.L., K.C., J.M., V.H. and X.T. analyzed data; X.T. wrote the paper.

## Acknowledgements

This work was supported by the SFARI Bridge to Independence grant, Charles Hood Foundation, SYNGAP Research Fund, PTEN Research fund, RSZ TNC fund, and NIH R01NS138758 awarded to X.T. This work was also supported by NINDS: R01NS105628 and R01NS102937, awarded to J.M. and supported E.C. and by 5R01MH104610 awarded to R.J. We graciously thank Dr. Sumeet Gupta from the Whitehead Institute genomics core facility, Dr. Liang Sun and Dr. Ashish Jain from the BCH Bioinformatics core facility for data analysis service. We thank Dr. Larry Benowitz for reading the paper and providing useful feedback.

## References

1. M. A. Lemmon, J. Schlessinger, Cell signaling by receptor tyrosine kinases. Cell 141, 1117–1134 (2010).

2. R. Kannaiyan, D. Mahadevan, A comprehensive review of protein kinase inhibitors for cancer therapy. Expert Rev Anticancer Ther 18, 1249–1270 (2018).

3. P. Tsapogas, C. J. Mooney, G. Brown, A. Rolink, The Cytokine Flt3-Ligand in Normal and Malignant Hematopoiesis. Int J Mol Sci 18, (2017).

4. J. Remnestal, L. Oijerstedt, A. Ullgren, J. Olofsson, S. Bergstrom, K. Kultima, M. Ingelsson, L. Kilander, M. Uhlen, A. Manberg, C. Graff, P. Nilsson, Altered levels of CSF proteins in patients with FTD, presymptomatic mutation carriers and non-carriers. Transl Neurodegener 9, 27 (2020).

5. J. U. Kazi, L. Ronnstrand, FMS-like Tyrosine Kinase 3/FLT3: From Basic Science to Clinical Implications. Physiol Rev 99, 1433–1466 (2019).

6. N. Daver, R. F. Schlenk, N. H. Russell, M. J. Levis, Targeting FLT3 mutations in AML: review of current knowledge and evidence. Leukemia 33, 299–312 (2019).

7. K. Mackarehtschian, J. D. Hardin, K. A. Moore, S. Boast, S. P. Goff, I. R. Lemischka, Targeted disruption of the flk2/flt3 gene leads to deficiencies in primitive hematopoietic progenitors. Immunity 3, 147–161 (1995).

8. O. deLapeyriere, P. Naquet, J. Planche, S. Marchetto, R. Rottapel, D. Gambarelli, O. Rosnet, D. Birnbaum, Expression of Flt3 tyrosine kinase receptor gene in mouse hematopoietic and nervous tissues. Differentiation 58, 351–359 (1995).

9. J. Ilzecka, Cerebrospinal fluid Flt3 ligand level in patients with amyotrophic lateral sclerosis. Acta Neurol Scand 114, 205–209 (2006).

10. C. Liao, V. Vuokila, H. Catoire, F. Akcimen, J. P. Ross, C. V. Bourassa, P. A. Dion, I. A. Meijer, G. A. Rouleau, Transcriptome-wide association study reveals increased neuronal FLT3 expression is associated with Tourette’s syndrome. Commun Biol 5, 289 (2022).

11. M. Dehlin, J. Bjersing, M. Erlandsson, N. Andreasen, H. Zetterberg, K. Mannerkorpi, M. Bokarewa, Cerebrospinal Flt3 ligand correlates to tau protein levels in primary Sjogren’s syndrome. Scand J Rheumatol 42, 394–399 (2013).

12. M. Shi, J. Bradner, A. M. Hancock, K. A. Chung, J. F. Quinn, E. R. Peskind, D. Galasko, J. Jankovic, C. P. Zabetian, H. M. Kim, J. B. Leverenz, T. J. Montine, C. Ginghina, U. J. Kang, K. C. Cain, Y. Wang, J. Aasly, D. Goldstein, J. Zhang, Cerebrospinal fluid biomarkers for Parkinson disease diagnosis and progression. Ann Neurol 69, 570–580 (2011).

13. W. Liedtke, Long March Toward Safe and Effective Analgesia by Enhancing Gene Expression of Kcc2: First Steps Taken. Front Mol Neurosci 15, 865600 (2022).

14. A. Tassou, M. Thouaye, D. Gilabert, A. Jouvenel, J. P. Leyris, C. Sonrier, L. Diouloufet, I. Mechaly, S. Mallie, J. Bertin, M. Chentouf, M. Neiveyans, M. Pugniere, P. Martineau, B. Robert, X. Capdevila, J. Valmier, C. Rivat, Activation of neuronal FLT3 promotes exaggerated sensorial and emotional pain-related behaviors facilitating the transition from acute to chronic pain. Prog Neurobiol 222, 102405 (2023).

15. C. Rivat, C. Sar, I. Mechaly, J. P. Leyris, L. Diouloufet, C. Sonrier, Y. Philipson, O. Lucas, S. Mallie, A. Jouvenel, A. Tassou, H. Haton, S. Venteo, J. P. Pin, E. Trinquet, F. Charrier-Savournin, A. Mezghrani, W. Joly, J. Mion, M. Schmitt, A. Pattyn, F. Marmigere, P. Sokoloff, P. Carroll, D. Rognan, J. Valmier, Inhibition of neuronal FLT3 receptor tyrosine kinase alleviates peripheral neuropathic pain in mice. Nat Commun 9, 1042 (2018).

16. W. Fiedler, S. Kayser, M. Kebenko, M. Janning, J. Krauter, M. Schittenhelm, K. Gotze, D. Weber, G. Gohring, V. Teleanu, F. Thol, M. Heuser, K. Dohner, A. Ganser, H. Dohner, R. F. Schlenk, A phase I/II study of sunitinib and intensive chemotherapy in patients over 60 years of age with acute myeloid leukaemia and activating FLT3 mutations. Br J Haematol 169, 694–700 (2015).

17. X. Tang, J. Drotar, K. Li, C. D. Clairmont, A. S. Brumm, A. J. Sullins, H. Wu, X. S. Liu, J. Wang, N. S. Gray, M. Sur, R. Jaenisch, Pharmacological enhancement of KCC2 gene expression exerts therapeutic effects on human Rett syndrome neurons and Mecp2 mutant mice. Sci Transl Med 11, (2019).

18. P. Q. Duy, M. He, Z. He, K. T. Kahle, Preclinical insights into therapeutic targeting of KCC2 for disorders of neuronal hyperexcitability. Expert Opin Ther Targets 24, 629–637 (2020).

19. M. M. Zack, R. Kobau, National and State Estimates of the Numbers of Adults and Children with Active Epilepsy - United States, 2015. MMWR Morb Mortal Wkly Rep 66, 821–825 (2017).

20. W. Loscher, H. Potschka, S. M. Sisodiya, A. Vezzani, Drug Resistance in Epilepsy: Clinical Impact, Potential Mechanisms, and New Innovative Treatment Options. Pharmacol Rev 72, 606–638 (2020).

21. M. Laplante, D. M. Sabatini, Regulation of mTORC1 and its impact on gene expression at a glance. J Cell Sci 126, 1713–1719 (2013).

22. P. W. Brownjohn, J. Smith, R. Solanki, E. Lohmann, H. Houlden, J. Hardy, S. Dietmann, F. J. Livesey, Functional Studies of Missense TREM2 Mutations in Human Stem Cell-Derived Microglia. Stem Cell Reports 10, 1294–1307 (2018).

23. P. Douvaras, B. Sun, M. Wang, I. Kruglikov, G. Lallos, M. Zimmer, C. Terrenoire, B. Zhang, S. Gandy, E. Schadt, D. O. Freytes, S. Noggle, V. Fossati, Directed Differentiation of Human Pluripotent Stem Cells to Microglia. Stem Cell Reports 8, 1516–1524 (2017).

24. A. Banerjee, R. V. Rikhye, V. Breton-Provencher, X. Tang, C. Li, K. Li, C. A. Runyan, Z. Fu, R. Jaenisch, M. Sur, Jointly reduced inhibition and excitation underlies circuit-wide changes in cortical processing in Rett syndrome. Proc Natl Acad Sci U S A 113, E7287–E7296 (2016).

25. S. Sivakumaran, R. A. Cardarelli, J. Maguire, M. R. Kelley, L. Silayeva, D. H. Morrow, J. Mukherjee, Y. E. Moore, R. J. Mather, M. E. Duggan, N. J. Brandon, J. Dunlop, S. Zicha, S. J. Moss, T. Z. Deeb, Selective inhibition of KCC2 leads to hyperexcitability and epileptiform discharges in hippocampal slices and in vivo. J Neurosci 35, 8291–8296 (2015).

26. D. S. Reddy, R. Kuruba, Experimental models of status epilepticus and neuronal injury for evaluation of therapeutic interventions. Int J Mol Sci 14, 18284–18318 (2013).

27. Y. Takagi, J. Harada, A. Chiarugi, M. A. Moskowitz, STAT1 is activated in neurons after ischemia and contributes to ischemic brain injury. J Cereb Blood Flow Metab 22, 1311–1318 (2002).

28. W. L. Hsu, Y. L. Ma, D. Y. Hsieh, Y. C. Liu, E. H. Lee, STAT1 negatively regulates spatial memory formation and mediates the memory-impairing effect of Abeta. Neuropsychopharmacology 39, 746–758 (2014).

29. J. Z. Tan, Y. Yan, X. X. Wang, Y. Jiang, H. E. Xu, EZH2: biology, disease, and structure-based drug discovery. Acta Pharmacol Sin 35, 161–174 (2014).

30. G. Ciceri, A. Baggiolini, H. S. Cho, M. Kshirsagar, S. Benito-Kwiecinski, R. M. Walsh, K. A. Aromolaran, A. J. Gonzalez-Hernandez, H. Munguba, S. Y. Koo, N. Xu, K. J. Sevilla, P. A. Goldstein, J. Levitz, C. S. Leslie, R. P. Koche, L. Studer, An epigenetic barrier sets the timing of human neuronal maturation. Nature 626, 881–890 (2024).

31. N. Khan, B. Schoenike, T. Basu, H. Grabenstatter, G. Rodriguez, C. Sindic, M. Johnson, E. Wallace, R. Maganti, R. Dingledine, A. Roopra, A systems approach identifies Enhancer of Zeste Homolog 2 (EZH2) as a protective factor in epilepsy. PLoS One 14, e0226733 (2019).

32. D. O’Carroll, S. Erhardt, M. Pagani, S. C. Barton, M. A. Surani, T. Jenuwein, The polycomb-group gene Ezh2 is required for early mouse development. Mol Cell Biol 21, 4330–4336 (2001).

33. M. Yeo, K. Berglund, G. Augustine, W. Liedtke, Novel repression of Kcc2 transcription by REST-RE-1 controls developmental switch in neuronal chloride. J Neurosci 29, 14652–14662 (2009).

34. L. Chen, L. Wan, Z. Wu, W. Ren, Y. Huang, B. Qian, Y. Wang, KCC2 downregulation facilitates epileptic seizures. Sci Rep 7, 156 (2017).

35. J. Tornberg, V. Voikar, H. Savilahti, H. Rauvala, M. S. Airaksinen, Behavioural phenotypes of hypomorphic KCC2-deficient mice. Eur J Neurosci 21, 1327–1337 (2005).

36. V. Magloire, J. Cornford, A. Lieb, D. M. Kullmann, I. Pavlov, KCC2 overexpression prevents the paradoxical seizure-promoting action of somatic inhibition. Nat Commun 10, 1225 (2019).

37. O. M. Pulido, Domoic acid toxicologic pathology: a review. Mar Drugs 6, 180–219 (2008).

38. A. R. Paciorkowski, R. N. Traylor, J. A. Rosenfeld, J. M. Hoover, C. J. Harris, S. Winter, Y. Lacassie, M. Bialer, A. N. Lamb, R. A. Schultz, E. Berry-Kravis, B. E. Porter, M. Falk, A. Venkat, R. J. Vanzo, J. S. Cohen, A. Fatemi, W. B. Dobyns, L. G. Shaffer, B. C. Ballif, E. D. Marsh, MEF2C Haploinsufficiency features consistent hyperkinesis, variable epilepsy, and has a role in dorsal and ventral neuronal developmental pathways. Neurogenetics 14, 99–111 (2013).

39. T. K. Creson, C. Rojas, E. Hwaun, T. Vaissiere, M. Kilinc, A. Jimenez-Gomez, J. L. Holder, Jr., J. Tang, L. L. Colgin, C. A. Miller, G. Rumbaugh, Re-expression of SynGAP protein in adulthood improves translatable measures of brain function and behavior. Elife 8, (2019).

40. Serena Tamura1, Andrew D. Nelson3,4†, Perry W.E. Spratt3,4†, Henry Kyoung3,4, Xujia Zhou1,2, Zizheng, Li1, Jingjing Zhao1,2, Stephanie S. Holden3,3,6, Atehsa Sahagun3,4, Caroline M. Keeshen3,4, Congyi Lu7, E.C.H., 4, Roy Ben-Shalom3,4, Jen Q. Pan7, J.T.P., 4,6, Stephan J. Sanders3,5; Navneet Matharu1, Nadav Ahituv1,2*, Kevin J. Bender3, 4*, CRISPR activation rescues abnormalities in SCN2A haploinsuffi ciency-associated autism spectrum disorder bioRxiv 10.1101/2022.03.30.486483, (2022).

41. I. E. Overwater, A. B. Rietman, A. M. van Eeghen, M. C. Y. de Wit, Everolimus for the treatment of refractory seizures associated with tuberous sclerosis complex (TSC): current perspectives. Ther Clin Risk Manag 15, 951–955 (2019).

42. Y. Li, H. Wang, J. Muffat, A. W. Cheng, D. A. Orlando, J. Loven, S. M. Kwok, D. A. Feldman, H. S. Bateup, Q. Gao, D. Hockemeyer, M. Mitalipova, C. A. Lewis, M. G. Vander Heiden, M. Sur, R. A. Young, R. Jaenisch, Global transcriptional and translational repression in human-embryonic-stem-cell-derived Rett syndrome neurons. Cell Stem Cell 13, 446–458 (2013).

43. X. Tang, L. Zhou, A. M. Wagner, M. C. Marchetto, A. R. Muotri, F. H. Gage, G. Chen, Astroglial cells regulate the developmental timeline of human neurons differentiated from induced pluripotent stem cells. Stem Cell Res 11, 743–757 (2013).

44. D. S. Svoboda, M. I. Barrasa, J. Shu, R. Rietjens, S. Zhang, M. Mitalipova, P. Berube, D. Fu, L. D. Shultz, G. W. Bell, R. Jaenisch, Human iPSC-derived microglia assume a primary microglia-like state after transplantation into the neonatal mouse brain. Proc Natl Acad Sci U S A 116, 25293–25303 (2019).

45. M. I. Love, W. Huber, S. Anders, Moderated estimation of fold change and dispersion for RNA-seq data with DESeq2. Genome Biol 15, 550 (2014).

46. M. Kanehisa, M. Furumichi, Y. Sato, M. Kawashima, M. Ishiguro-Watanabe, KEGG for taxonomy-based analysis of pathways and genomes. Nucleic Acids Res 51, D587–D592 (2023).

47. C. Gene Ontology, S. A. Aleksander, J. Balhoff, S. Carbon, J. M. Cherry, H. J. Drabkin, D. Ebert, M. Feuermann, P. Gaudet, N. L. Harris, D. P. Hill, R. Lee, H. Mi, S. Moxon, C. J. Mungall, A. Muruganugan, T. Mushayahama, P. W. Sternberg, P. D. Thomas, K. Van Auken, J. Ramsey, D. A. Siegele, R. L. Chisholm, P. Fey, M. C. Aspromonte, M. V. Nugnes, F. Quaglia, S. Tosatto, M. Giglio, S. Nadendla, G. Antonazzo, H. Attrill, G. Dos Santos, S. Marygold, V. Strelets, C. J. Tabone, J. Thurmond, P. Zhou, S. H. Ahmed, P. Asanitthong, D. Luna Buitrago, M. N. Erdol, M. C. Gage, M. Ali Kadhum, K. Y. C. Li, M. Long, A. Michalak, A. Pesala, A. Pritazahra, S. C. C. Saverimuttu, R. Su, K. E. Thurlow, R. C. Lovering, C. Logie, S. Oliferenko, J. Blake, K. Christie, L. Corbani, M. E. Dolan, H. J. Drabkin, D. P. Hill, L. Ni, D. Sitnikov, C. Smith, A. Cuzick, J. Seager, L. Cooper, J. Elser, P. Jaiswal, P. Gupta, P. Jaiswal, S. Naithani, M. Lera-Ramirez, K. Rutherford, V. Wood, J. L. De Pons, M. R. Dwinell, G. T. Hayman, M. L. Kaldunski, A. E. Kwitek, S. J. F. Laulederkind, M. A. Tutaj, M. Vedi, S. J. Wang, P. D’Eustachio, L. Aimo, K. Axelsen, A. Bridge, N. Hyka-Nouspikel, A. Morgat, S. A. Aleksander, J. M. Cherry, S. R. Engel, K. Karra, S. R. Miyasato, R. S. Nash, M. S. Skrzypek, S. Weng, E. D. Wong, E. Bakker, T. Z. Berardini, L. Reiser, A. Auchincloss, K. Axelsen, G. Argoud-Puy, M. C. Blatter, E. Boutet, L. Breuza, A. Bridge, C. Casals-Casas, E. Coudert, A. Estreicher, M. Livia Famiglietti, M. Feuermann, A. Gos, N. Gruaz-Gumowski, C. Hulo, N. Hyka-Nouspikel, F. Jungo, P. Le Mercier, D. Lieberherr, P. Masson, A. Morgat, I. Pedruzzi, L. Pourcel, S. Poux, C. Rivoire, S. Sundaram, A. Bateman, E. Bowler-Barnett, A. J. H. Bye, P. Denny, A. Ignatchenko, R. Ishtiaq, A. Lock, Y. Lussi, M. Magrane, M. J. Martin, S. Orchard, P. Raposo, E. Speretta, N. Tyagi, K. Warner, R. Zaru, A. D. Diehl, R. Lee, J. Chan, S. Diamantakis, D. Raciti, M. Zarowiecki, M. Fisher, C. James-Zorn, V. Ponferrada, A. Zorn, S. Ramachandran, L. Ruzicka, M. Westerfield, The Gene Ontology knowledgebase in 2023. Genetics 224, (2023).

48. N. E. Sanjana, O. Shalem, F. Zhang, Improved vectors and genome-wide libraries for CRISPR screening. Nat Methods 11, 783–784 (2014).

49. F. A. Ran, P. D. Hsu, J. Wright, V. Agarwala, D. A. Scott, F. Zhang, Genome engineering using the CRISPR-Cas9 system. Nat Protoc 8, 2281–2308 (2013).

